# Parameter tuning is a key part of dimensionality reduction via deep variational autoencoders for single cell RNA transcriptomics

**DOI:** 10.1101/385534

**Authors:** Qiwen Hu, Casey S. Greene

## Abstract

Single-cell RNA sequencing (scRNA-seq) is a powerful tool to profile the transcriptomes of a large number of individual cells at a high resolution. These data usually contain measurements of gene expression for many genes in thousands or tens of thousands of cells, though some datasets now reach the million-cell mark. Projecting high-dimensional scRNA-seq data into a low dimensional space aids downstream analysis and data visualization. Many recent preprints accomplish this using variational autoencoders (VAE), generative models that learn underlying structure of data by compress it into a constrained, low dimensional space. The low dimensional spaces generated by VAEs have revealed complex patterns and novel biological signals from large-scale gene expression data and drug response predictions. Here, we evaluate a simple VAE approach for gene expression data, Tybalt, by training and measuring its performance on sets of simulated scRNA-seq data. We find a number of counter-intuitive performance features: i.e., deeper neural networks can struggle when datasets contain more observations under some parameter configurations. We show that these methods are highly sensitive to parameter tuning: when tuned, the performance of the Tybalt model, which was not optimized for scRNA-seq data, outperforms other popular dimension reduction approaches – PCA, ZIFA, UMAP and t-SNE. On the other hand, without tuning performance can also be remarkably poor on the same data. Our results should discourage authors and reviewers from relying on self-reported performance comparisons to evaluate the relative value of contributions in this area at this time. Instead, we recommend that attempts to compare or benchmark autoencoder methods for scRNA-seq data be performed by disinterested third parties or by methods developers only on unseen benchmark data that are provided to all participants simultaneously because the potential for performance differences due to unequal parameter tuning is so high.

## 1. Introduction

Single-cell RNA sequencing (scRNA-seq) profiles the transcriptomes of individual cells [1], allowing researchers to study heterogeneous cell characteristics and responses [2, 3]. Due to the small amount of RNA captured in each cell as well as technical factors related to capture efficiency, scRNA-seq data have a high dropout rate (many genes have no measured expression in each cell). Researchers often analyze these data by projecting cells into a low dimensional space, which enables downstream analysis such as imputation of missing measurements and visualization.

Widely used approaches include the linear principal component analysis (PCA) [4], which doesn’t take dropout into account, and ZIFA [5], which uses zero-inflated factor analysis to model the dropout events and do dimension reduction. The t-distributed stochastic neighbor embedding (t-SNE) method is also widely used [6]. This method uses local structure, but it is time consuming for large datasets and has been reported to be highly sensitive to hyperparameters [7]. The recently proposed Uniform Manifold Approximation and Projection (UMAP) [8] method attempts to address these limitations by preserving more global structure and as much local structure as t-SNE. These approaches do not model the dropout characteristic of scRNA-seq data.

Deep generative neural network models can learn low-dimensional representations from large amounts of unlabeled data and have been successfully applied to many domains, such as image and text generation [9]. Variational autoencoders (VAE) learn this representation by compressing data into a constrained, low-dimensional space [10]. VAEs have been used in biology to analyze large-scale gene expression data and drug response predictions [11, 10]. In recent months, preprints proposing numerous deep neural network models for scRNA-seq data have been posted. Grønbech et al. [12] proposed a Gaussian-mixture VAE model for raw counts from scRNA-seq data and found the model can learn biologically groupings of scRNA-seq dataset. Eraslan et al. developed a deep count autoencoder based on zero-inflated negative binomial noise model for data imputation [13]. Lopez et al. developed single-cell Variational Inference (scVI) based on hierarchical Bayesian models, which can be used for batch correction, dimension reduction and identification of differentially expressed genes [14]. Deng et al. propose an autoencoder that includes a feedback step after zeroes are imputed [15]. These methods often report performance, but while many report hyperparameter selections, few describe how those parameters were reached.

In this work, our goal was to understand the extent to which reported performance of the neural network methods was due modifications for scRNA-seq data. We applied a straightforward VAE developed for bulk gene expression data, Tybalt [10], to simulated and real scRNA-seq data under various parameter settings. Some performance characteristics, including a decrease in performance when the number of examples was increased, suggest substantial sensitivity to hyperparameters. We sought to optimize parameters and adjust the dimensionality of the model to rescue performance. In our prior work from *PSB 2015* using autoencoders for the analysis of bulk gene expression data, performance was relatively stable over many parameter values [16]. In contrast, the performance of the standard VAE, Tybalt, changes from dismal to better than other popular dimension reduction approaches – PCA, ZIFA, UMAP and t-SNE – with only modest parameter tuning.

These results should guide the reporting of new methods. First, it is critically important that reviewers expect manuscripts in this area to report the extent to which hyperparameters affect performance across multiple datasets. Second, manuscripts reporting new techniques should be evaluated both on theoretical grounding as well as empirical results. Because results can be changed easily by light tuning, self-reported performance numbers may provide only weak evidence. Third, assessments and benchmarking should be done by disinterested parties with a realistic amount of parameter tuning or should be performed by first parties on datasets for which the labels are not revealed until after predictions are made.

## 2. Methods

### 2.1. Data Simulation

We simulated scRNA-seq data using Splatter [17]. We used the default simulation parameters provided by Splatter to generate synthetic scRNA-seq data with variable numbers of genes, cell types, cells, outliers, etc. We simulated data with variable numbers of cells (ncell: 500 - 5000), genes (nGenes: 20000 - 60000), cell types (nGroups: 5 – 15) and probabilities of expression outliers (outlier: 0.1 – 0.5). In total, we generated 40 simulated single-cell datasets. We normalized the raw count matrix by TPM (Transcripts Per Kilobase Million).

### 2.2. Model Structure and Training

VAEs model the distribution P(X) of data in a high dimensional space from a low dimensional latent space z. VAEs consist of two connected neural networks: the encoder and decoder. Data are compressed by the encoder and reconstructed by the decoder. The variation probability Q(z|X) is used to approximate the posterior distribution P(z|X), which is then optimized to minimize the Kullback–Leibler (KL) divergence between Q(z|X) and P(z|X) and reconstruction loss [18, 19]. A baseline model for gene expression data, termed Tybalt and which we use here, was described in [10]. The encoder was a multi-layer (varied from 0 to 2) neural network. The representative latent space z was sampled from a Gaussian distribution q_θ_(*z*|*X*), with mean and variance generated by the encoder network. The learned latent space z was used to re-generate the count matrix X’ by the decoder, which was also a multi-layer neural network (from 0 to 2) (Figure 1). For the first stage, we trained Tybalt with three structures: a one-layer model with a gene-wise TPM vector connected to 20 latent features and then reconstructed output; a two-layer model with the TPM vector encoded into a 100-node hidden layer, then the 20 latent features, then a 100-node hidden layer, and then the reconstructed output; and a three-layer model which contains two 100-node hidden layers. The model was built in Keras (version 2.0.6) [20] with a TensorFlow (Version 1.0.1) backend [21].

**Figure 1:**
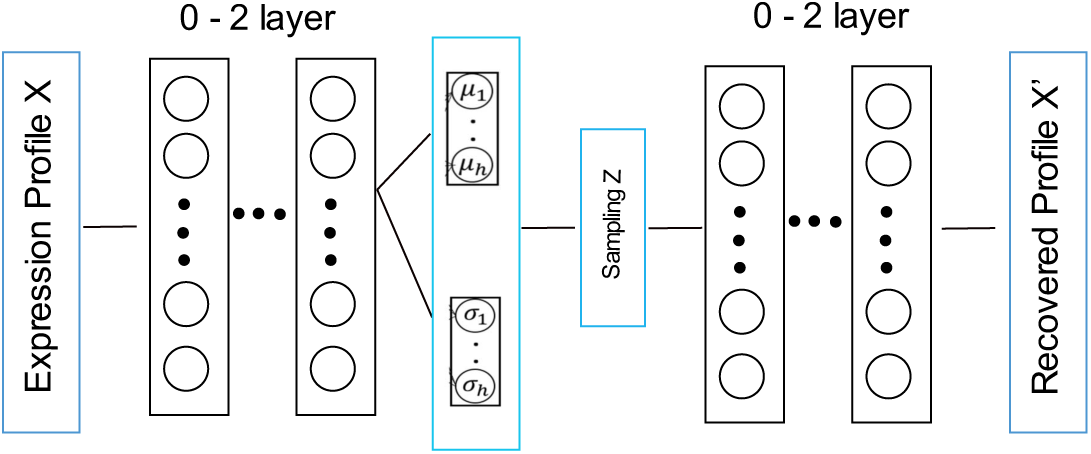
Overview of the structure of variation autoencoder. The model consists an encoder network and a decoder network, both of them are designed as 0-2 layers fully connected neural networks.

### 2.3. Parameter Tuning

We tuned parameters using a grid search over batch size (50, 100, 200), epochs (25, 50, 100, 200), neural network depth (2, 3) and, for models with two or more layers, the dimensionality of the first layer (100, 250, 500). Simulated data were partitioned into training and test data, with the test set being 10% of the full data. For real data, we selected three single cell datasets with author-assigned cell type labels [25 - 27]. We downloaded count matrices from the Hemberg Group repository of data (https://hemberg-lab.github.io/scRNA.seq.datasets/). We zero-one normalized the count matrix before training the VAE.

### 2.4. Performance Measurement

We used three evaluation metrics to measure performance: 1) *k*-means based 2) *k*NN based 3) average silhouette score. The *k*-means based and *k*NN based measurements measure how well the low-dimensional space allows simple methods to recover simulated cell types. The average silhouette score measures the extent to which clusters are separable in the latent space. An ideal method is accurate and produces separable clusters.

#### 2.4.1. k-means performance assessment

In the *k*-means based evaluation we performed iterative *k*-means clustering on the low-dimensional latent space. We compared the predicted clustering results with the known cell types in the simulated data. We performed *k*-means clustering for 50 times to get a stable measurement and – to evaluate a best-case scenario – we set the number of clusters, *k*, to the number of true cell types in the data. We assessed methods by the normalized mutual information (NMI) [22], between cell types and the known categories as well as the adjusted rand index (ARI) [23].

#### 2.4.2. kNN performance assessment

For the *k*NN evaluation, we used *k* nearest neighbors to predict cell type from latent space distances and assessed performance by 5-fold cross validation within the simulated dataset. To more closely replicate how methods are used in practice, the model was tuned within only the training data by a sweep over the neighbor number parameter with 3-fold cross validation. We assessed performance using accuracy, precision, recall and f-score, but report only accuracy due to space constriants.

#### 2.4.3. Average silhouette score performance assessment

We used the silhouette score [24] to measure the extent to which simulated cell type clusters are internally close in the latent space but separated from other cell types. The silhouette value is between −1 to 1. A silhouette value of 1 indicates that the data point is of distance zero from other points of the same type, while one of −1 indicates that the point is distance zero from all points of a different cluster but some distance from points of the same cluster. Average silhouette score over all points then indicates how separable each cell type is in the latent space.

## 3. Results

### 3.1. The performance of multiple methods on simulated data

We tested the performance of five dimension reduction approaches: Tybalt, ZIFA, UMAP, t-SNE and PCA using the three different evaluation metrics over the data simulated by Splatter [17] as described in the Methods section. We selected metrics that would be sensitive to the quality of reduced representations (k-means, knn, and silhouette width) because our goal was to assess these representations and not to build the best possible cell type predictor. The k-means clustering approach evaluates the extent to which a hypersphere in the latent space is capable of capturing cell types accurately. We used both NMI and ARI to measure performance, though results for each are relatively similar so we present only NMI within the main text due to space constraints. The kNN approach evaluates the extent to which local structure in the latent space reflects cell type. The silhouette width approach evaluates the extent to which the within-type distances in the latent space are smaller than the between-type distances.

#### 3.1.1. k-means based results

The performance of most methods varied substantially under simulation parameters (Figure 2). As expected, more cell types led to reduced performance, assessed via NMI, of PCA, ZIFA, and the variational autoencoders. As the number of cells changed, the performance of ZIFA and PCA fluctuated. Intriguingly, the three-layer VAE, which had the most parameters to fit and which should have improved with more data, performed worse as the number of cells increased. Later we show that this result is due to substantial parameter sensitivity. Less surprisingly, increasing the number of genes (and consequently parameters) reduced the performance of larger autoencoders. Outliers reduced the performance of PCA but had relatively inconsistent effects on other methods. For the default parameters, the two-layer Tybalt model was generally high-performing, but both the one- and three-layer models showed variable performance. This surprising sensitivity to simulated data characteristics suggests that VAEs may be very sensitive the fit between parameters and data.

**Figure 2:**
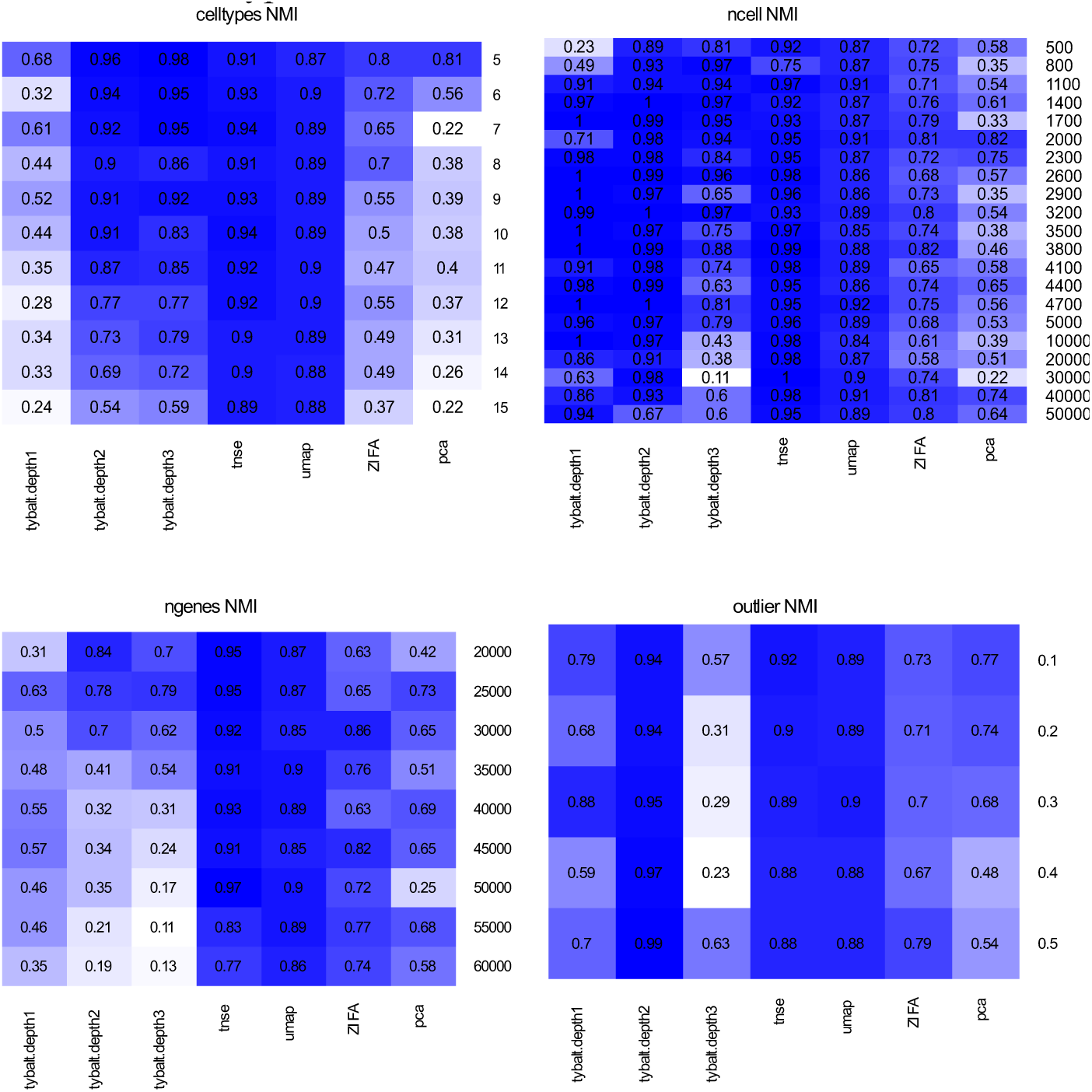
Performance of different dimensionality reduction approaches based on simulated single cell datasets measured by normalized mutual information (NMI)

#### 3.1.2. kNN and silhouette score results

Results based on the kNN and silhouette evaluations are consistent with the results from k-means. We display results for representative datasets to show variability. The GitHub repository contains complete results. Performing kNN in the latent space revealed relatively poor performance of the linear methods (Figure 3, PCA and ZIFA). UMAP and t-SNE perform well across many combinations, and the VAEs generally performed reasonably well until the number of genes became very high, presumably because the number of parameters leads to insufficient training data.

**Figure 3:**
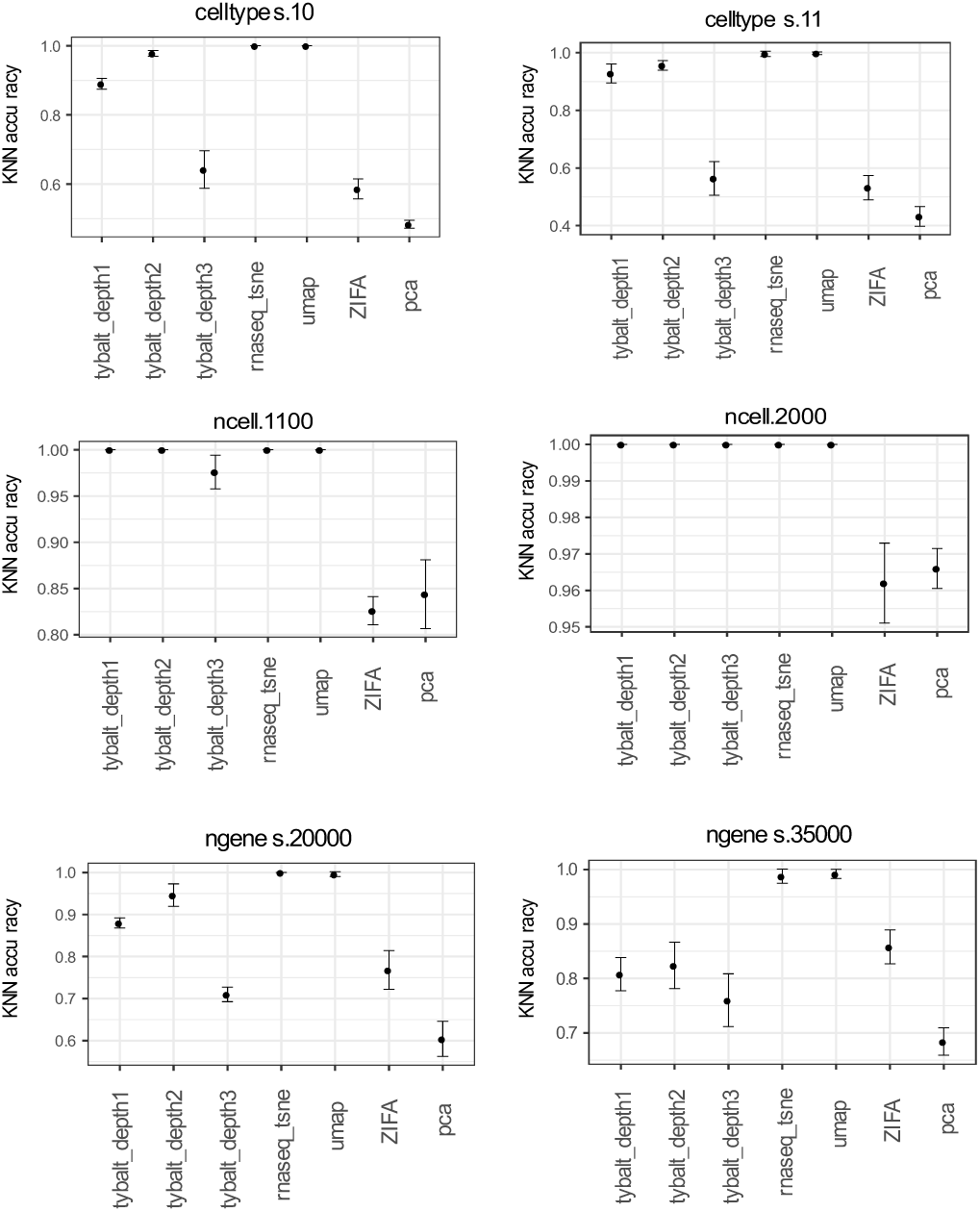
kNN performance for representative simulated single cell datasets under different parameters. Error bars show the standard deviation of accuracy across cross validation intervals, and stability differed between methods.

The silhouette score evaluation tests something slightly different than the k-means and kNN evaluations. While those focus on the extent to which there is some detectable separation between cell types, the silhouette score evaluates the extent to which within cell-type distances are smaller than between cell-type distances. Despite this difference, the results remain consistent (Figure 4) with the other evaluations. As the number of cell types increases, the performance of all method is drops, though the decrease is somewhat less pronounced with t-SNE and UMAP. This evaluation also shows the same unexpected performance drop as the number of cells (thus, examples) increases with three-layer VAE models.

**Figure 4:**
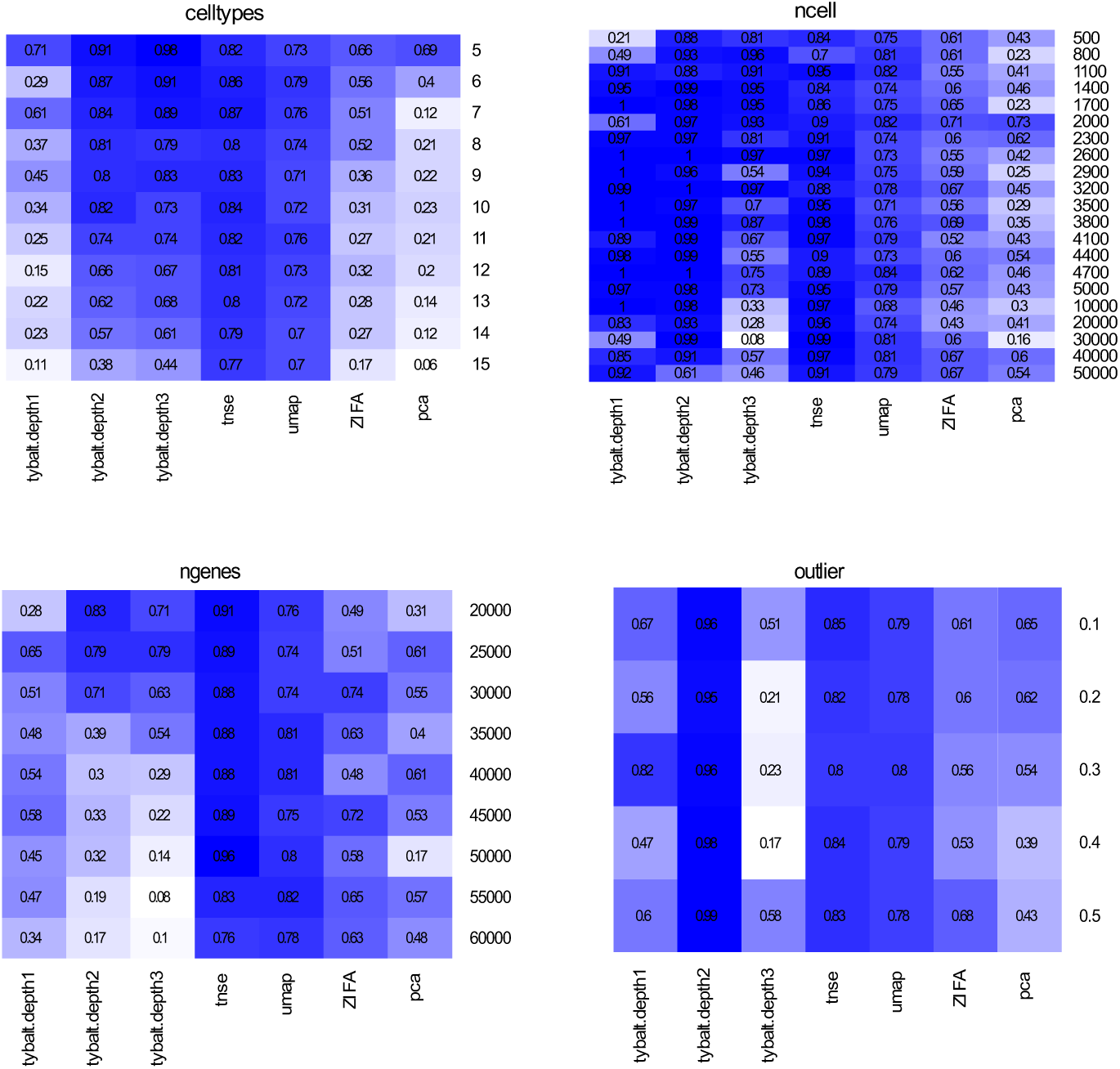
Performance of different dimensionality reduction approaches based on average silhouette score.

#### 3.1.3. Summary of the performance comparison

Our results indicate that no dimensionality reduction method outperformed the rest in all cases. The performances of the linear methods (PCA and ZIFA) were generally poorer under the cases that we tested, and the performances of t-SNE and UMAP were generally quite robust within the bounds that were tested. Perhaps the most interesting finding of this stage of the analysis was that the VAE-based methods struggled in expected situations (i.e., when the number of genes, and consequently parameters, increased) but also in unexpected situations (i.e., when the number of cells, and consequently training examples, increased). This suggested that either the model structure or parameter combinations must be poor, because otherwise more examples would always lead to better performance. We explore the implications of this finding more fully in Section 3.3.

### 3.2. 2-dimensional projection of simulated datasets

To visualize the results associated with the evaluation described in 3.1, we projected cells into the learned latent spaces and then reduced those spaces to 2-dimensional space via t-SNE on the latent space values while coloring by the simulated cell types (Figure 5). We observe performance characteristics that hint at why the methods exhibited strong or poor performance in different settings. For example, with few outliers the structure of t-SNE remains reasonable, but as the number of outliers increases some points begin to shift to the extremes of the projection. UMAP generally has high between-group distances and low within-group distances and is not affected by cell types. The linear methods (ZIFA and PCA) along with the single-layer variational autoencoder (Tybalt) appear to unequally space the cell types, even though these are not correlated with each other. The two- and three-layer Tybalt models do not have this relationship, though the three-layer model appears to train poorly with more outliers.

**Figure 5:**
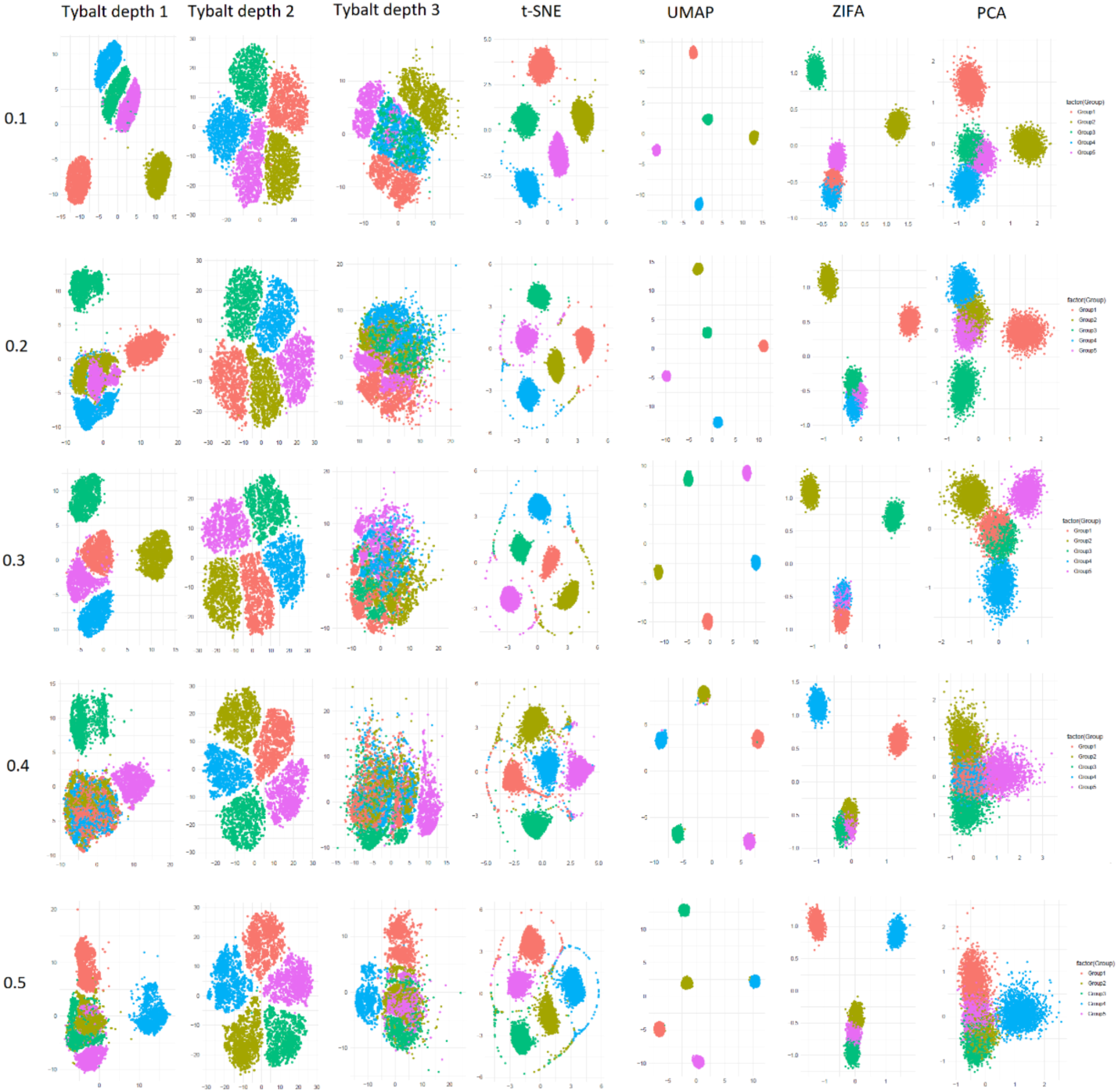
2-dimensional projection of simulated single cell data using one-, two- and three-layer VAE models (Tybalt), t-SNE, ZIFA, UMAP and PCA based on different proportions of outliers (0.1 – 0.5).

### 3.3. Analysis of VAE performance failures

As observed in the previous section, we found that the three-layer Tybalt model’s performance dropped precipitously under certain conditions. Our hypothesis was that the hyperparameters were not appropriate for this setting. We sought to determine the extent to which we could rescue performance under the least expected failure mode from Section 3.1: namely that performance dropped when the number of examples increased. We performed a parameter sweep as described in section 2.3. Note that this grid search is of a very modest size, so we would expect modest performance changes. Results for the k-means evaluation are shown in Table 1. We noticed that the performance of VAE changes dramatically during parameter selection. In this case, performance varies from dismal to better than most the other dimensionality reduction approaches. With 30,000 cells the worst three-layer model has an NMI of zero, while the best has an NMI of 0.96 (Table 1).

**Table 1.** Best and worst parameter values for two- and three-layer Tybalt models with many cells for simulated datasets. l: learning rate, b: batch size, e: epoch, c: dimensionality of the first hidden layer.

| **2-layer model** |  |  |  |  |  |  |  |  |  |  |  |  |
| --- | --- | --- | --- | --- | --- | --- | --- | --- | --- | --- | --- | --- |
| Best combination |  |  |  |  |  |  | Worst combination |  |  |  |  |  |
| ncells | l | b | e | c | NMI | ARI | l | b | e | c | NMI | ARI |
| 10000 | 0.0005 | 200 | 25 | 100 | 0.99 | 0.99 | 0.0005 | 50 | 200 | 500 | 0.08 | 0.05 |
| 20000 | 0.001 | 200 | 25 | 250 | 0.98 | 0.95 | 0.001 | 200 | 200 | 500 | 0.2 | 0.13 |
| 30000 | 0.0005 | 100 | 100 | 100 | 0.97 | 0.97 | 0.0005 | 50 | 100 | 500 | 0 | 0 |
| **3-layer model** |  |  |  |  |  |  |  |  |  |  |  |  |
| Best combination |  |  |  |  |  |  | Worst combination |  |  |  |  |  |
| ncells | l | b | e | c | NMI | ARI | l | b | e | c | NMI | ARI |
| 10000 | 0.001 | 50 | 100 | 100 | 1 | 1 | 0.002 | 200 | 200 | 500 | 0.06 | 0.04 |
| 20000 | 0.0005 | 100 | 25 | 250 | 0.97 | 0.95 | 0.001 | 100 | 100 | 500 | 0.02 | 0.01 |
| 30000 | 0.0005 | 100 | 25 | 250 | 0.96 | 0.94 | 0.0005 | 50 | 200 | 250 | 0 | 0 |

We also selected three single cell datasets of various cell numbers and tissues where author-assigned sample labels were available. Baron et al. [25] and Wang et al. [26] assay the human pancreas with 8569 and 635 cells respectively. Camp et al. [27] measured 777 cells from human liver tissue. As with simulated data, VAE performance changed substantially after parameter tuning, although the range of reasonable parameters appears to be broader than in simulated data (Table 2).

**Table 2.** Best and worst parameter values for two- and three-layer Tybalt models for real datasets. l: learning rate, b: batch size, e: epoch, c: dimensionality of the first hidden layer.

| **2-layer model** |  |  |  |  |  |  |  |  |  |  |  |  |
| --- | --- | --- | --- | --- | --- | --- | --- | --- | --- | --- | --- | --- |
| Best combination |  |  |  |  |  |  | Worst combination |  |  |  |  |  |
| Datasets | l | b | e | c | NMI | ARI | l | b | e | c | NMI | ARI |
| Baron et al. | 0.0005 | 100 | 25 | 100 | 0.64 | 0.38 | 0.002 | 50 | 200 | 500 | 0.36 | 0.17 |
| Wang et al. | 0.001 | 200 | 200 | 500 | 0.46 | 0.3 | 0.0005 | 200 | 25 | 100 | 0.2 | 0.11 |
| Camp et al. | 0.002 | 50 | 25 | 500 | 0.81 | 0.71 | 0.0005 | 100 | 25 | 100 | 0.64 | 0.47 |
| **3-layer model** |  |  |  |  |  |  |  |  |  |  |  |  |
| Best combination |  |  |  |  |  |  | Worst combination |  |  |  |  |  |
| Datasets | l | b | e | c | NMI | ARI | l | b | e | c | NMI | ARI |
| Baron et al. | 0.0005 | 100 | 25 | 500 | 0.63 | 0.36 | 0.001 | 100 | 200 | 500 | 0.33 | 0.17 |
| Wang et al. | 0.0005 | 50 | 200 | 500 | 0.45 | 0.3 | 0.0005 | 200 | 25 | 100 | 0.24 | 0.13 |
| Camp et al. | 0.0005 | 200 | 50 | 500 | 0.76 | 0.62 | 0.0005 | 200 | 25 | 250 | 0.61 | 0.42 |

We projected cells into the latent space learned by Tybalt models pre- and post-tuning to visualize the effect of hyperparameters (Figure 6). The two-layer model was robust within the tested range. With the optimal parameters there was a slightly larger gap between cell types, but the cell types were still clearly separated. For the three-layer model there were substantial differences. Before tuning, the three-layer model shows some signs of a failure to train, which could explain the poor quantitative performance. After tuning, the cell types were clearly separated. These results demonstrate that parameter tuning dramatically affects performance for VAE models in this domain. In the case we evaluated, this appears to be more pronounced with the deeper neural network. However, it is also possible that the default parameters that we selected to tune around happened to be a relatively robust space for two-layer networks for scRNA-seq data.

**Figure 6:**
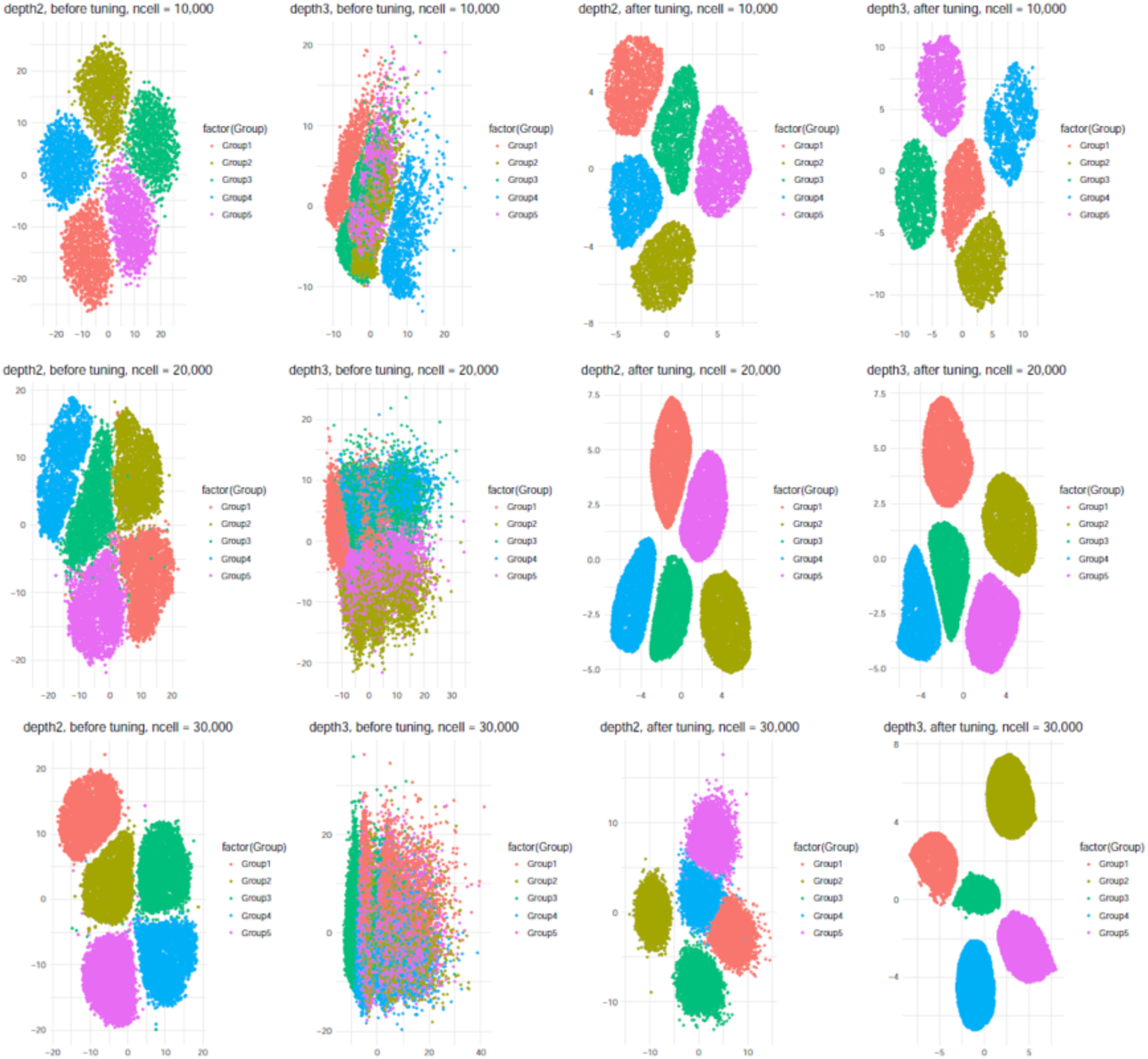
Parameter tuning improve the performance for deeper network. 2-dimensional projection of simulated single cell data using tybalt_depth2 and Tybalt_depth3 before and after parameter tuning.

## 4. Conclusions

Certain preprints now report good performance for deep neural network methods using VAEs or other types of autoencoders for the analysis of scRNA-seq data [12, 13, 14, 15]. In certain cases, the authors report performance using a set of parameters (see Table 2 of Lopez et al.) without reporting how hyperparameters were tuned or how performance varied through tuning [14]. This poses a particular challenge when authors report performance comparisons with other methods. For example, Deng et al. [15] compare their scScope method with scVI, but they report “we followed the same parameter setting in the original study Lopez et al. [14] and setting the latent dimension to 50.” However, Lopez et al., report numerous potential parameter combinations, so which ones were used is impossible to interpret.

We sought to understand the extent to which reported variability in performance was due to differences in methods versus differences in parameter settings. Thus, we evaluated the performance of a simple VAE model developed for bulk gene expression data, Tybalt, under various parameter settings. We find that, in many cases, a base VAE of two layers performs similarly to other methods. However, we also find substantial performance differences with hyperparameter tuning. Though this is not entirely unexpected, the sensitivity of this class of methods under various parameter settings is not widely reported in the literature. Indeed, papers sometimes neglect to report the extent to which parameters were tuned and the extent to which authors optimize the parameter settings of other methods is unclear.

Wolpert and Macready [28] reported a No Free Lunch theorem that states that improved performance of an optimizer on one problem is paired with a decrease in performance in some other area. Our results suggest that algorithms that are more sensitive to parameters, combined with a publication process that encourages method developers to compare their own approaches to others, may experience a Continental Breakfast Included (CBI) effect. We term this the CBI effect because it accrues primarily to certain methods in specific settings. The CBI effect arises when researchers expend more researcher degrees of freedom [29] on their own method instead of other methods. The CBI effect is particularly strong when methods are highly sensitive to parameters, because the results change more substantially with each researcher degree of freedom that is expended.

Our results indicate evaluation of model performance based on empirical results can be misleading in the presence of the CBI effect. For example, we are able to make performance on the same dataset for a three-layer neural network vary from near random to near perfect (Section 3.3). At the current time, we recommend that authors who which to apply these methods expect to perform parameter tuning to achieve acceptable performance, which is likely to require substantially more compute time than is often reported because many manuscripts report only the compute time to train the final model. Moving forward, an unbiased approach is important for model evaluation and comparison. We recommend that authors developing these methods refrain from emphasizing comparisons unless methods are equally tuned and/or some sort of blinded design is used to control researcher degrees of freedom. It may be most practical to rely primarily on disinterested third parties or challenge-based frameworks for comparisons between methods.

## 5. Reproducibility

We provide the source code and scripts to reproduce the analysis at https://github.com/greenelab/CZI-Latent-Assessment/tree/master/single_cell_analysis

